# Physiochemically distinct SU-8 surfaces tailor *Xylella fastidiosa* cell-surface holdfast and colonization

**DOI:** 10.1101/2021.12.14.472636

**Authors:** Silambarasan Anbumani, Aldeliane M. da Silva, Andrei Alaferdov, Marcos V. Puydinger dos Santos, Isis G. B. Carvalho, Mariana de Souza e Silva, Stanislav Moshkalev, Hernandes F. Carvalho, Alessandra A. de Souza, Monica A. Cotta

## Abstract

SU-8 polymer is an excellent platform for diverse applications due to its high aspect ratio of micro/nanostructures fabrication and exceptional optical, chemical, and biocompatible properties. Although SU-8 has been often investigated for a variety of biological applications, how its surface properties influence both the interaction of bacterial cells with the substrate and its colonization is poorly understood. In this work, we tailor SU-8 nanoscale surface properties to investigate single cell motility, adhesion and successive colonization of a phytopathogenic bacteria, *Xylella fastidiosa*. Different surface properties of SU-8 thin films have been prepared using photolithography processing and oxygen plasma treatment. We found a significant difference in bacterial cell behavior and subsequent colonization on SU-8 as surface property changes. A larger density of carboxyl groups in hydrophilic plasma-treated SU-8 surfaces promotes faster cell motility in the earlier stage of the growth. The hydrophobic nature of pristine SU-8 surfaces has no trackable bacterial motility with 5 to 10 times more single cells adhered to surface than its plasma-treated counterpart. In fact, plasma-treated SU-8 samples suppressed bacterial adhesion, with surfaces showing less than 5% coverage. These results not only showcase that SU-8 surface properties can impact the bacterial behavior in a spatiotemporal manner, but also provide insights on the prominent ability of pathogens to evolve and adapt to different surface properties.

## 1. Introduction

The development of portable, high-throughput and cost-effective healthcare and medical devices is indebted to substantial advances in materials. The SU-8 epoxy polymer is one such material that has great potential for fabrication of high aspect ratio of micro/nanostructured scaffolds for lab-on-a-chip devices[1–4]. In particular, SU-8 has been used as an impressive platform for the development of various smart biomedical devices including biosensors[5–7], bacterial diagnosis[8–10], cantilever[11], bioelectrodes[12] and microrobots[13,14] owing to its excellent optical and mechanical properties with chemical stability even against acids[3,15,16]. Moreover, SU-8 nanostructures with tunable high aspect ratios have been fabricated and used for eukaryote cell interface and for simultaneous visualization of the traction force and focal adhesion of eukaryotic cells[17]. Whilst the SU-8 properties combined with its biocompatibility facilitates its application in bio-related material interfaces[6–9,17,18], how the surface properties of SU-8 affect bacterial adhesion and successive proliferation is poorly understood.

The evolution of bacteria relies on several survival strategies and as a collective response of multicellular assembly[19,20]. Specifically, the virulence of a pathogenic bacteria depends on its capability to attach to biotic and abiotic surfaces [21–23]. Interaction between bacteria and material surfaces can be affected by various properties of the material surface, including wettability (hydrophobic or hydrophilic), surface energy and chemistry, charge, elastic modulus, topography, and so on [21,24–27]. An in-depth understanding of the physicochemical aspects of bacterial-substratum interface is important both fundamentally and clinically for preventing microbial adhesion and consequently, biofilm infections [20,26–29]. Moreover, the development of *in vitro* models to unraveling the spatiotemporal relationship to bacterial adhesion, cell motility and subsequent colonization has remained a key topic in biomedical research[22,30].

Here, we investigate the *in vitro* cell motility, adhesion, and biofilm formation in the early-stage bacterial life cycle of *Xylella fastidiosa* with the different surface properties of SU-8. *X. fastidiosa* is a gram-negative phytopathogen that causes diseases worldwide in important crops (e.g. citrus, grape, coffee, almond, olives, among others); the bacteria colonizes two distinct habitats: xylem vessels of host plants, and the foregut of xylem-feeding insects which are transmission vectors [31]. In particular, the entire biofilm formation process, starting from single cell adhesion, has been investigated, turning *X. fastidiosa* into a reliable bacterial model[24,32]. This species is also interesting as it shares many genetic traits with other human bacteria [33,34] and has relatively slow duplication time (~6h) [30], which renders easier the observation of surface colonization. Moreover, *X. fastidiosa* relies on type-IV pili, which are about 2-to-6-μm long, for twitching motility; these pili are significantly impacted by surface chemistry[35,36]. The twitching motility governed by type IV pili is used to colonize different surfaces and play a crucial role in the development of biofilms of *X. fastidiosa* [35,36] as well as of several other Gram-negative bacteria [37–39].

In this work, different wettability and surface chemical modification of SU-8 samples have been achieved using UV illumination, and oxygen plasma treatment, to explore the characteristics of bacterial-SU-8 surface interfaces. Atomic Force Microscopy (AFM), contact angle measurements and X-Ray Photoelectron Spectroscopy (XPS) were used to evaluate nanoscale surface properties. The bacteria-surface interaction and its relation to the nanoscale surface properties of the samples is investigated by monitoring cell motility, adhesion and microcolonies formation using confocal laser scanning microscopy (CLSM) for different time intervals (6, 12, and 24 hours) of growth, under identical environments. The mean velocity, net displacement and mean square displacement (MSD) of single cell trajectories have been extracted from cell tracking information for different samples. We found that carboxylic functional surfaces resulted in straighter trajectories with faster-moving cells, for shorter times of observation. The hydrophobic surface of pristine SU-8 leads to relatively absence of cell movement while a larger number of cells adhered to surface. Furthermore, the influence of physiochemically distinct surfaces in the development of microcolonies of the bacteria has been studied.

## 2. Experimental section

### 2.1. Sample preparation

SU-8 monomer films deposited by spin coating are exposed to UV illumination so that a polymer film of thickness of about 300 nm was obtained. For all bacterial adhesion experiments carried out in this work, the samples were cleaned to remove inorganic as well as organic contamination. For the preparation of oxygen plasma-treated samples, a radio frequency (RF) plasma generator (PLASMA Technology SE80, Model ACG 5) operating at frequency of 13.56 MHz and base pressure of 1 mTorr was used with the following treatment conditions. During treatment, O_2_ gas flow of 50 sccm at a pressure of approximately 100 mTorr and plasma power of 100 W was maintained for treatment times of 60 seconds in O_2_ flow. The detailed optimization of the treatment time is given elsewhere [5]. In addition to SU-8 samples, flat InP substrates are also used for comparison. The substrates were cleaned with acetone, isopropanol, and deionized water, and dried with a gentle nitrogen flow. The InP substrates were further cleaned by oxygen plasma using O_2_ gas flow of 50 sccm at a pressure of approximately 100 mTorr and plasma power of 200 W was maintained for 10 minutes in O_2_ flow. To generate carboxylic-acid-modified InP surfaces, initially the oxygen plasma cleaned samples has been incubated in DMSO (dimethyl sulfoxide) containing 5M Ethanolamine hydrochloride for overnight. Then the samples were washing using DI water and further PEGylated by depositing 2 mM of amino-reactive, heterobifunctional NHS-PEG-COOH (MW 3.400, LaysanBio, USA)[40]. After the functionalization process, the supports were rinsed three times in water and dried with N_2_ gas. For all bacterial growth experiments, the substrates were sterilized by UV lamp in the biosafety hood (VECO Biosafe A1) immediately prior to the experiment.

### 2.2. Surface characterization

#### 2.2.1. Surface Topography

Keysight (model 5500) scanning atomic force microscopy (AFM) with intermittent contact mode (AC mode) using the cantilever of a nominal stiffness of 3 N/m was used to record the surface topography of samples. Root mean square (RMS) surface roughness of the samples was extracted from the topography images with 0.7 × 0.7 μm^2^ area using Gwydion (version 2.56) software.

#### 2.2.2. Contact angle

Static water contact angle of each surface was measured using sessile drop contact angle measurement method[5,41]. For each sample, three measurements were performed at ambient temperature of approximately (25 ± 2) °C and relative humidity ranging from 50% to 70%. For the samples incubated in PW media, the surface contact angle was determined after brief rinse with DI water followed by drying with a gentle nitrogen flow.

#### 2.2.3. XPS measurement

The analysis of surface chemical groups was performed using X-ray photoelectron spectroscopy (XPS). XPS experiments were carried out on a SPECS system (SPECS GmbH, Germany) equipped XR 50 - X-ray source with Al Kα radiation (*hν*= 1486.6 eV) and Phoibos 100 hemispherical energy analyzer with MCD-9 detector. The X-ray anode was run at 100 W and the high voltage was kept at 10.0 kV with sample normal polar angle of tilt of 20°. The pass energy was fixed at 20.0 eV to ensure sufficient sensitivity. The base pressure of the analyzer chamber was about 3 × 10^-10^ mbar.

The quantitative analysis was performed in program CasaXPS (Version 2.3.16Dev52). The C_1s_ lines were investigated in detail. In order to establish the type of chemical bond between carbon and oxygen atoms, deconvolution of these lines was performed using the program PeakFit v 4.06 (PeakFit^TM^). The data were fitted by Voigt functions using method “I-residuals” for detection of hidden peaks after performing a Shirley background subtraction.

### 2.3. Bacterial growth

Green fluorescent protein (GFP) expressing strain 11399 of *Xylella fastidiosa* subsp. *pauca* bacteria was used in this study[42]. Bacterial inoculum with a concentration of 1×10^7^ CFU/mL (OD_600_ 0.5) from the pre-inoculum was used for the experiments as initial concentration for bacterial growth studies in PW broth media[43]. Bacteria extraction and pre-inoculum preparation was described elsewhere[32,42]. The SU-8 and InP samples were incubated for different growth times (specified in the respective manuscript text) in a bacterial stove (410/3NDR, Nova Ética, Brazil) at 28°C without culture media replacement.

### 2.4. Image acquisition using Confocal microscopy

For *in-vitro* studies, the samples were placed inside a Teflon dish (10 mm diameter and 5 mm in height). Then, 400 μL of inoculum was injected inside the Teflon dish, which was covered with a sterilized borosilicate cover glass. The assembly was incubated inside a bacterial oven at 28°C. The bacterial growth on the various samples were studied using confocal laser scanning microscope (CLSM) (Carl Zeiss AG, Germany) operated with 40x water-immersion objective (Plan-Apochromat, NA. 1.0, Zeiss). Each sample has been studied at 6, 12, 24 hours of growth (the samples were stored back inside a bacterial oven at 28 °C for more hours after each observation). The exposure times and lamp intensity were kept optimal to reduce photobleaching and phototoxicity. *X. fastidiosa* cell motility on the surface were recorded using time-lapse movies acquired in every two second interval for 1 to 5 minutes. For quantification of cell attachment and colonization to the surfaces, 2D images were collected at separate fields of view of 210 × 210 μm^2^ for each sample.

### 2.5. Image analysis

The area of bacterial biofilms was extracted from raw fluorescence images, and 3D rendered images were created using Carl Zeiss Zen (3.0 SR) software. The image backgrounds were subtracted using the built-in rolling-ball background subtraction algorithm in ImageJ software. The cell movement trajectories were tracked on these different samples using tracking algorithm provided in Track mate plugin in ImageJ according to the methodology described by Tinevez *et.al* [44,45]. Laplacian of Gaussian filter was applied to detect single cells, and estimated blob diameter and threshold were fixed for the optimum detection of cells in each data collection.

Net track displacement values, which represent the length of a straight line that extends from the first to the last spot of a track, and mean velocity values, which provides the average of the velocity between successive spots over all the spots of a track, have been extracted from the cell tracking information[44] of the CLSM images. The angle between successive (x,y) coordinates (segments) of the tracks is calculated to investigate the orientation of the tracks. Furthermore, to explore positional freedom which the bacteria experience, mean square displacement (MSD) as a function of time has been calculated from Isy software using the track information[46]. The MSD describes the average of the squared distances between the starting and ending position of a cell movement for all timelags (Δt) within a trajectory.

### 2.6. Statistics

All data are presented as mean value ± standard deviation. The statistical significance between different groups has been accessed by resulting from two-tailed, unpaired t-tests. A *p*-value smaller than 0.05 was considered as statistically significant. Number of asterisks on figures indicate data statistical significance level (\**p* < 0.05; \*\**p* < 0.01; \*\*\**p* < 0.001; n.s. = non-significant).

## 3. Results and Discussion

Despite the SU-8 surface being inherently hydrophobic, which is undesired for most biological applications, the modification of surface properties such as wettability and surface functional groups can be employed by means of dry and wet treatment processes [5,7,47,48]. To evaluate the surface properties, we chose SU-8 pristine and oxygen plasma treated (for 60 seconds) SU-8 samples, based on our optimized protocol for surface modifications that provides minimal surface roughness[5]. In addition, flat Indium Phosphide (InP) substrates are also chosen for reference control since the *X. fastidiosa* bacterial adhesion to InP surfaces has been well studied[49,50]. Initially, RMS surface roughness (**Fig. 1A**) was characterized from AFM topography images (**Fig. S1**), and contact angle measurements were used to study surface wettability of the InP, pristine and plasma-treated SU-8 samples (**Fig. 1B**). The results show that pristine SU-8 and InP samples have relatively small surface roughness (< 0.5 nm) and a more hydrophobic nature (contact angle > 65□). On the other hand, plasma treatment of SU-8 surfaces provided an increase in the surface roughness features (~1.5 nm) and a highly hydrophilic nature as compared to the pristine surface.

**Figure 1.**
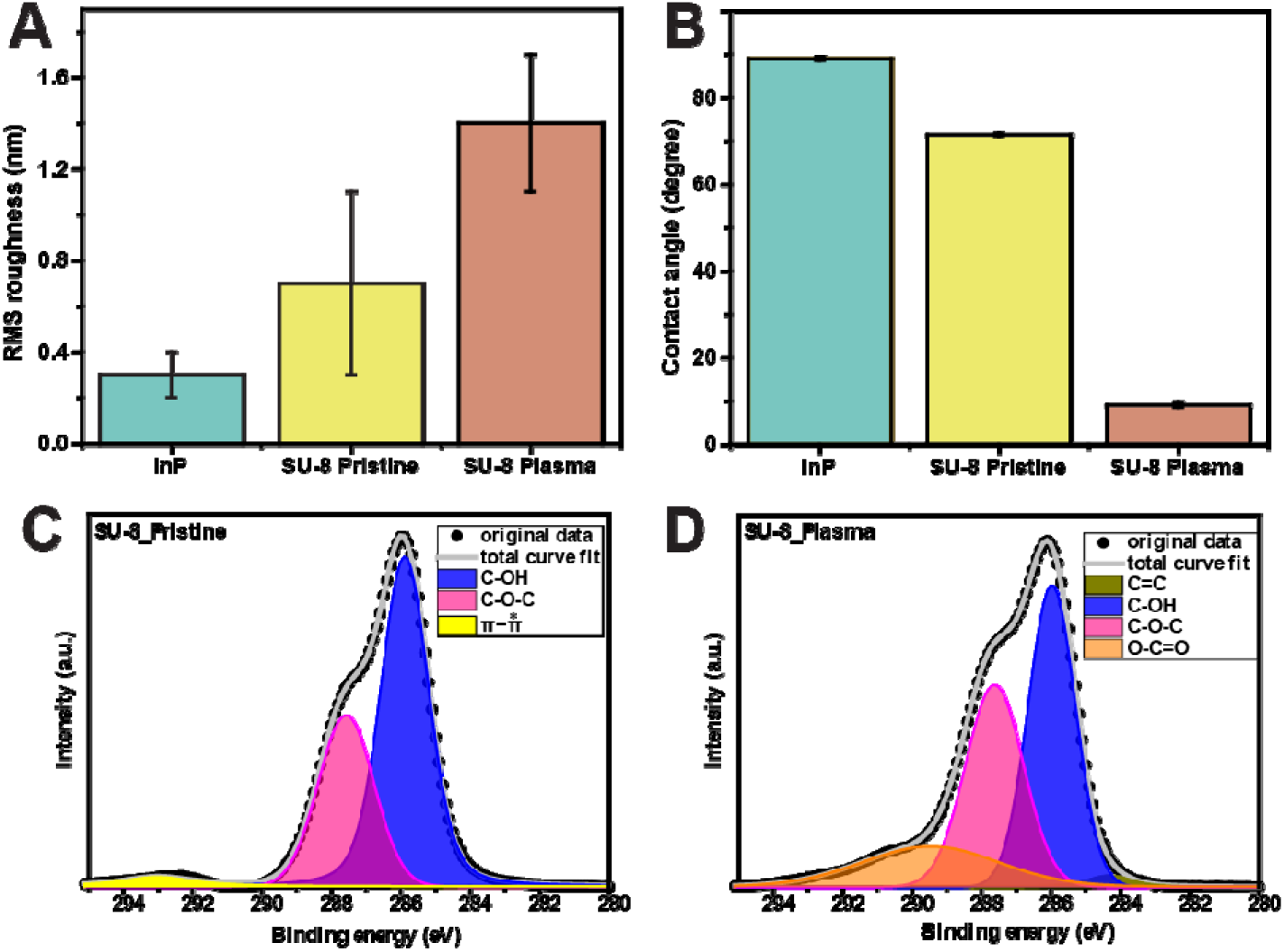
Surface physicochemical characterization of SU-8 polymer. (A) RMS roughness measurements extracted from AFM topography images and (B) contact angle of InP, SU-8 pristine and SU-8 plasma-treated samples. Deconvoluted C_1s_ peaks of XPS spectrum of (C) pristine and (D) plasma-treated samples.

The surface functional groups of the SU-8 samples were characterized by XPS (**Fig. 1C and D**). Using data represented in literature [51–58] the few components of C_1s_ peak found after deconvolution procedure can be assigned. Components located at 283.6-283.9 eV and 284.3 eV are associated with non-oxygenated carbon bounded atoms in pentagon and heptagon rings (C-C) bonded and sp^2^ carbon (C=C) conformations. Components at 285.4-286.0 eV, 287.0 - 287.6 eV and 288.3 eV are attributed to C-atoms bonded to O in ethers group (C-OH), C in epoxy groups (C-O-C), and carbon bonded to two oxygen atoms in carbonyl groups (C=O), respectively. It should be noted that the plasma-treated samples showed a component at 288.4-289.5 eV that represents carboxylate (O-C=O) groups. Moreover, one broad component was detected at a high-energy region (peak near 292-293 eV) usually associated with π-π* electrons transition processes[59,60]. A comparison of the components areas which are proportional to the amount of the chemical bonds is shown in Table 1 for pristine and plasma treated SU-8 samples. These results quantitatively confirm that pristine samples present surface epoxy groups. In the case of plasma-treated samples, an additional surface carboxylic group was observed[5].

**Table:1.** Comparison of the C_1s_ peak components area of pristine and plasma-treated SU-8 surfaces obtained from XPS spectra.

| Surface | Area of $C_{1s}$ peak components (in %) | | | | |
| --- | --- | --- | --- | --- | --- |
| | C-C | C-O-C | C-OH | O-C=O | $\pi-\pi^*$ peak |
| SU-8 pristine | absent | 34.5 | 62.2 | absent | 3.3 |
| SU-8 plasma | 1.9 | 36.5 | 44 | 17.6 | absent |

In order to investigate the surface wetting properties of the samples, the contact angle has been measured after incubating the samples in PW growth media (**Table 2**) that we used later for the bacterial growth. The pristine SU-8 surfaces remained hydrophobic until 6 h of incubation. For longer incubation times (12 and 24 h) the surface becomes moderately hydrophilic. Interestingly, the plasma-treated surface still maintains its hydrophilic nature even after 24 h incubation time, while InP surfaces show moderate hydrophilic behavior for all three different incubated times. The SU-8 samples of distinct surface wettability (hydrophobic and hydrophilic) properties with the enhanced surface carboxylic groups have been extended next to investigate the *X. fastidiosa* bacterial growth experiments along with comparison to the InP samples.

**Table 2.** Contact angle measurements of SU-8 and InP surfaces, after incubation in PW media for 6, 12 and 24 h.

| Surface | Contact angle after incubating the samples in PW media for |  |  |
| --- | --- | --- | --- |
|  | 6 h | 12 h | 24 h |
| **InP** | $53 \pm 0.5$ | $48.8 \pm 0.9$ | $49 \pm 2.0$ |
| **SU-8 pristine** | $81.5 \pm 1.9$ | $51.3 \pm 1.6$ | $53 \pm 0.5$ |
| **SU-8 plasma treated** | $24.4 \pm 1.5$ | $23.7 \pm 0.8$ | $38.8 \pm 1.3$ |

*X. fastidiosa* bacterial growth experiments have been studied using CLSM for the interval of 6, 12 and 24h growth time. Initially, we monitored the bacterial motility from time-lapse videos by tracking the motion of each cell on InP, SU-8 pristine and plasma-treated surfaces prepared according to protocols mentioned in the methods section. **Fig. 2 (A-C)** shows trajectories (in yellow) of motile single cells after 6 hours of growth. Plasma-treated samples have comparatively straighter and longer moving cell trajectories up to 6 h (**Fig. 2B**). However, for longer growth times (12 and 24 h) of these samples and all growth times of pristine InP samples, trajectories presented random orientations and curly type, associated with twitching motility (**Fig. 2A** & **Fig. S2**). Surprisingly, the hydrophobic pristine SU-8 samples show absolutely no trackable moving cells; instead, more adherent single cells than the plasma treated SU-8 and InP samples are present, even after a prolonged growth time (24h) (**Fig. 2C**). The cells irreversibly adhered to pristine SU-8 surfaces within the first contact (**Video S1**); strongly adhered cells make it difficult to capture the cell trajectory on the surface for the observation time windows used here.

**Figure 2.**
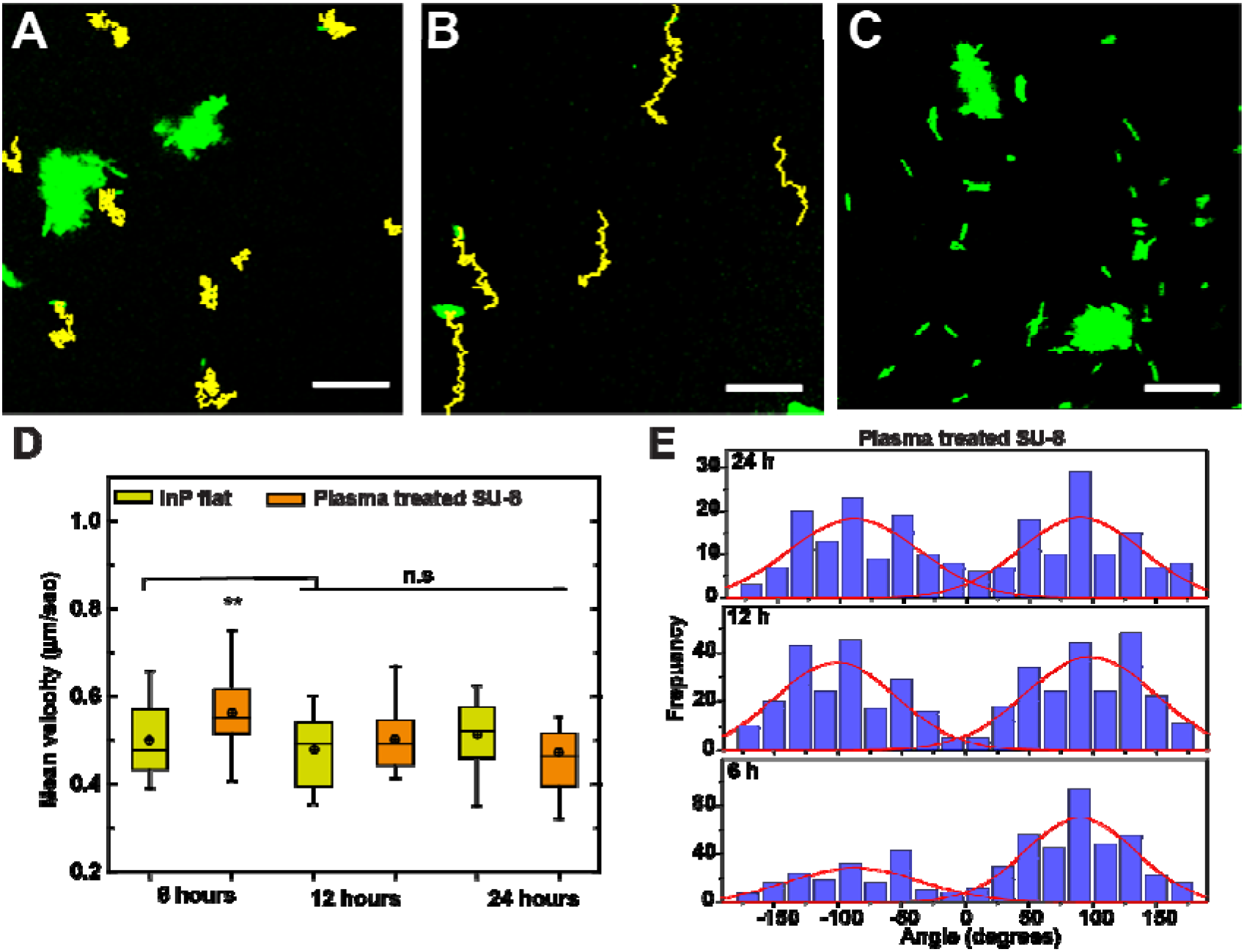
*X. fastidiosa* single cell motility changes on different surface physicochemical properties of samples. Trajectory of moving *X. fastidiosa* single cells on (A) InP (B) plasma treated SU-8 and (C) pristine SU-8 (no trackable motile cells) from time lapse CLSM videos for 3 min tracking after 6 h of growth. Yellow lines indicate cell trajectory. Scale bars: 20 μm. (D) Mean velocity of single cell movement on hydrophilic InP and plasma treated SU-8 surfaces after 6, 12 and 24 h growth (from N=25 tracks for each). (E) Distribution of angle of each segment of the trajectories obtained from xy coordinates of the tracks on plasma-treated SU-8 samples after 6, 12 and 24 h growth; red lines indicate Gaussian fitting to positive and negative ranges of angle distribution.

Mean velocity and orientation of cell trajectories is acquired from cell tracking information. The comparison of continuous time intervals of 1, 3 and 5 minutes and different locations (**Fig. S3**) on plasma-treated SU-8 samples after 6 h of growth, confirm no significant differences in the mean velocity (~0.5 μm/s) associated with the time interval and different location of the measurements. Therefore, mean velocity and displacement values for 3 min of tracking events have been extracted and analyzed by statistical comparison. After 6 hours of growth, the average cell motility on plasma-treated SU-8 surfaces is significantly higher (~0.56 μm/s) than the mean velocity of later grown samples (12 and 24h) and all three-growth times of InP surfaces as well (~0.5 μm/s) (**Fig. 2D**). The average cell velocity of about 0.5 μm/sec (~30 μm/min) of the cells moving on these surfaces is comparable with previously reported mean velocity of *X. fastidiosa* (~25 μm/min) studied under no-flow condition in microfluidic chambers[35]. The individual segments of each trajectory were calculated from the xy coordinates extracted from tracking information for the plasma-treated surface (**Fig. 2E**). Positive angles are two times larger than negative after 6h growth, which confirms the straighter orientation of the cells (**Fig. 2E** & **Table S1**), while 12 and 24 h samples have almost the same movement for both ranges of the angle distribution histograms.

Moreover, cell trajectories in plasma treated SU-8 surfaces after 6 h of growth (**Fig. 3A**) showed significantly longer displacement values (~15 μm) than all other cases (6, 12 and 24 h) of growth on InP and, 12 and 24h growth on plasma-treated SU-8 samples (~7 μm). The distinct straighter orientations, larger mean velocity and displacement of trajectories of plasma-treated SU-8 after 6h growth could be due to the larger density of surface carboxyl groups, and the hydrophilic nature of the sample; since plasma was applied just before inoculation, the activated carboxyl groups may have a role in cell motility. However, after 12 and 24h growth, the motility drops on the plasma-treated samples, most likely due to surface conditioning by both growth medium and continuous EPS (Extracellular Polymeric Substance) secretion[32] which covers the surface and, consequently, stabilize the carboxylic groups. To further confirm whether the carboxyl functional group is critical for such exquisite cell movement, InP samples functionalized with carboxylic groups (as explained in Methods section) were studied. Interestingly and in agreement with our hypothesis, the single cells trajectories on InP carboxylic surface are also straighter, with longer displacements for the 6 h growth period (**Fig. S4**).

**Figure 3.**
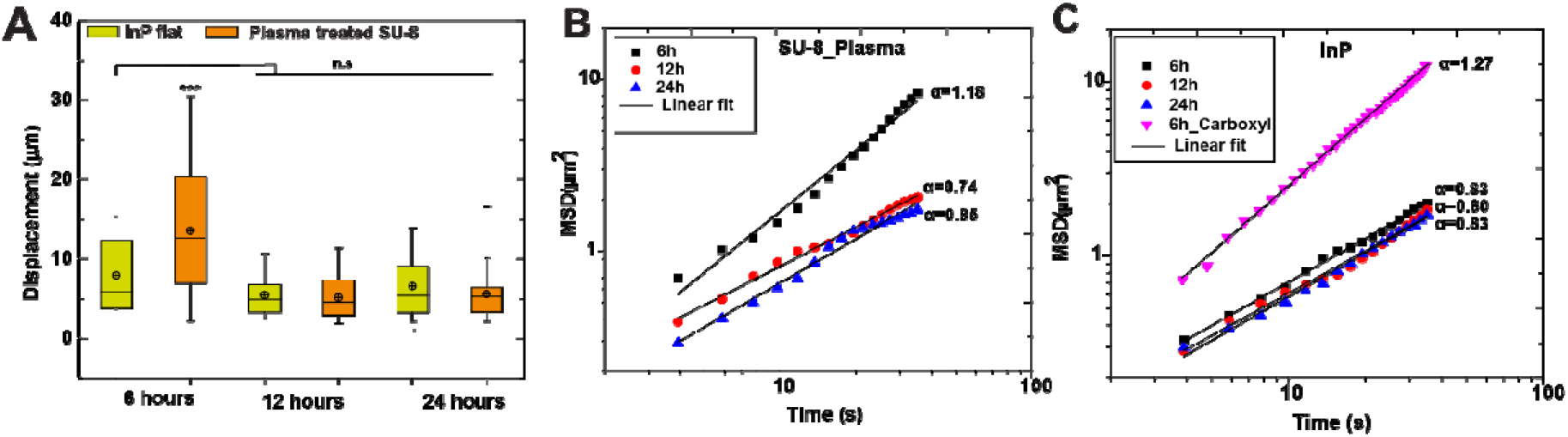
Quantifying displacement indicators of single cell trajectories on different surfaces. (A) Net displacement values for single cell movement on InP and plasma treated SU-8 surfaces after 6, 12 and 24 h growth (from N=25 tracks for each). (B) Average of mean square displacement (MSD) with function of time of single cell movement on plasma treated SU-8 and (C) InP surfaces after 6, 12 and 24 h of growth along with and carboxylic group functionalized InP after 6h growth; Black line indicates linear fit and α is the corresponding slope value (from N=10 tracks for each).

We further investigate the possible displacement of single cells in a more sensitive way by calculating averaged MSD as a function of time for all three-growth times of plasma-treated and InP samples, as shown in **Fig. 3B and C**, respectively. The log-log plot of MSD describes the anomalous diffusion model and the slope of the plot can indicate the characteristics of a cell trajectory[61]. For random diffusion, the MSD is expected to show slope α = 1, while slope =2 is related to ballistic motion[62]. Therefore, cell trajectories with MSD slopes α>1 (superdiffusive process) indicate persistent movement and slopes α <1 (subdiffusive process) suggest confined movement[61,63,64]. Indeed, the MSD curves of the plasma-treated samples indicate two distinct types of trajectories observed on the samples: persistent (slope=1.18 for 6 h growth: **Fig. 3B**) and confined (slope ≈ 0.8 for 12 & 24 h growth; **Fig 3B**). On the other hand, the MSD values for InP samples for all growth times indicate confined motion (slope ≈ 0.8; **Fig. 3C**) except for the carboxylated InP surface for 6h growth which indicates persistent motion (slope=1.27; **Fig. 3C**). Moreover, both plasma-treated SU-8 and carboxylated InP samples for 6 h growth time present MSD values one order of magnitude larger than their counterparts in Figures 3B and 3C, respectively.

Twitching motility occurs by pili elongation, surface adhesion and retraction[35,65], with type IV pili of *X. fastidiosa* typically found at one of the cell poles. The pili are composed of pilin proteins in which the pilus core is filled by amino-terminal α-helices and the outer surface is formed by carboxyl terminal domains[37]. In the case of carboxylic functional surfaces, a repulsive molecular interaction between *X. fastidiosa* pili and surface carboxylic groups most likely makes the surface less resistant against motion, also providing a longer distance exploration along the trajectory. Particularly, the larger slope of MSD curves (α>1) for the carboxylated samples until 6 h growth time would indicate the relatively smooth surface detachment and successive re-attachment of pili. Conrad *et al.* distinguish walking and crawling motions of *Pseudomonas aeruginosa* based on the MSD plots[63]. Regarding *X. fastidiosa* twitching motility, a previous study exhibit cell trajectories oriented towards upstream flow under fluidic conditions while random orientation of trajectories has been observed under no-fluidic conditions[35]. However, our study shows that surface chemistry can modify the bacterial behavior, as we confirmed the characteristics of persistent (carboxylated surface until 6 h) and confined (all other surfaces) crawling motion of *X. fastidiosa* present in our SU-8 samples, under no-flow conditions.

Since the distinct characteristics of bacterial motility are observed with different surfaces and growth times, we further compared how the number of single cells changed with respect to the surface properties of the samples. Single cells on the surface were counted from confocal images for each 6, 12, and 24 h growth (**Fig. 4 A, B**). The results show InP surface has increasing number of single cells over 6 and 12 h growth; after 24h growth, the number of cells remains almost stable. In contrast, single cells on the plasma-treated SU-8 surfaces have gradually decreased over time (6-24h). This could be due to the lack of surface interactions and larger displacement of the cells in the initial growth time on the plasma treated SU-8 sample. In other words, the cells are vastly motile to explore the desired surface, but that ultimately leads to dispersal of cells on surface due to weaker cellsurface interactions. In fact, the increased hydrophilicity of plasma-treated samples should reduce the number of adhered bacteria compared to pristine SU-8 as per general thermodynamic theory[66]. Our results confirmed this prediction, as we observed about 5-times larger number of individual attached cells to the pristine SU-8 surfaces after 6, 12 and 24h growth (**Fig. 4B**). Furthermore, more firmly-adhered single cells were observed for the pristine samples as indicated by arrows in the figure, showing single cells in the same surface location after 12 and 24 h of growth (**Fig. 4C)**. Also, the fast adhesion (**Supplementary Video S1)** indicates the strongest cell interaction with pristine SU-8 surfaces. Such scenario could be more likely due to type IV pili interaction to the hydrophobic nature of such surfaces, and further facilitated by the strong interaction of hydrophobic *X. fastidiosa* cell surface[66,67]. This strong adhesion of type IV pili to hydrophobic surfaces is also reported for *Pseudomonas aeruginosa*[68]. Moreover, our observations agree qualitatively with previous results on Gram-negative *Escherichia coli* adherence to polymeric surfaces with contact angle of 90° [69] and also with Gram-positive bacteria *Micromonospora purpurea* onto material surfaces with contact angle ranging from 54 to 130° [70]. It is interesting to note that the present observation on the stronger and fast adhesion to pristine SU-8 surfaces provides some support to the previous hypothesis that *X. fastidiosa* bacterial cells acquired from plants are quickly adhesive once they contact the hydrophobic-in-nature cuticle surface of foreguts [71,72].

**Figure 4.**
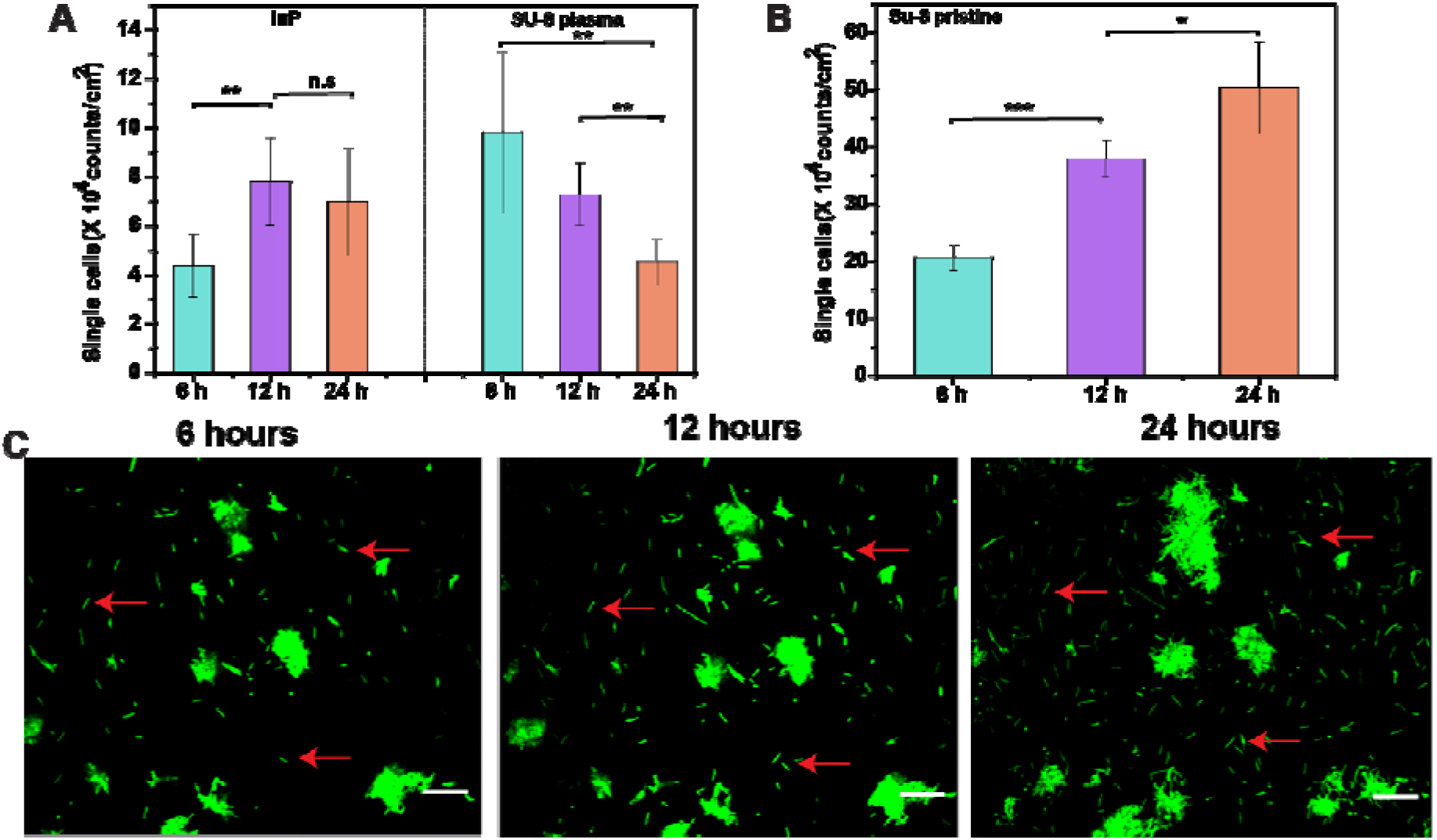
Bacterial adhesion on different surface properties (A) Number of single cells that exhibit motion or remain adhered on InP and plasma-treated SU-8 surfaces after 24h growth (from N=7 images each). (B) Number of single cells adhered on pristine SU-8 surfaces after 6, 12 and 24h growth (from N=7 images each). (C) *In vitro* CLSM images of bacterial adhesion on same area of a pristine SU-8 sample after 6, 12 and 24h growth. Red arrows point the few single cells firmly adhered to the same location of the surface, along the whole period of growth. Color histogram is adjusted so that single cells can be visualized; Scale bar:20 μm.

To determine how SU-8 surface properties influence bacterial growth, we next sought to quantify the microcolonies sizes and coverage area on the three different samples from the recorded *in vitro* CLSM images (**Fig. 4C and Supplementary Fig. S5).** The results confirm that larger microcolonies sizes occur on InP surfaces (**Fig. 5A**) for all growth times (6-24h), when compared to the other two surfaces. Especially, the higher bacterial coverage of *X. fastidiosa* after 24h growth (**Fig. 5B**) could be due to the large stiffness of the rigid InP surface; such a high colonization rate on rigid surfaces has been previously observed for several Gram-negative bacteria, including *X. fastidiosa* [73,74]. On the other hand, gradual growth of microcolonies, reaching up relatively large sizes over time, were observed for pristine SU-8 samples despite no trackable motile cells even after 24h growth (**Fig. 5A**). This observation is also in agreement with previous reports stating that hydrophobic interactions are usually the strongest of all long-range noncovalent interactions and they play a significant role in cell adhesion and aggregation[75–77]. In contrast, the plasma-treated sample present microcolonies with smaller sizes (**Fig. 5A**) and no significant change in size for all growth times used here. Furthermore, the coverage area is significantly smaller (70-90 %) when compared to pristine SU-8 and InP samples (**Fig. 5B**).

**Figure 5.**
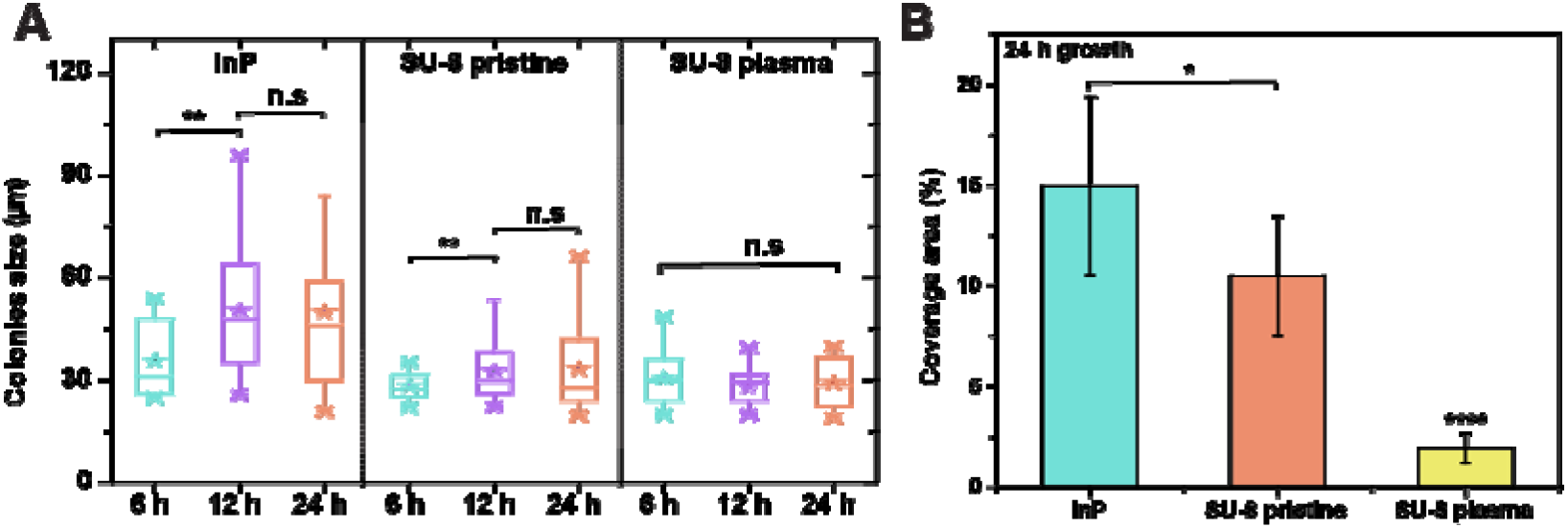
Bacterial colonization on distinct surface properties of samples. Comparison of (A) of microcolonies formed after 6, 12 and 24 h growth and (B) bacterial coverage percentage on the InP, pristine and plasma treated SU-8 surfaces after 24h growth. (N=10 images for each).

The interaction of bacterial cells with surface carboxylic groups and hydrophilic surface of plasma-treated surface affect both the attachment rate and colonization of bacteria, and the surface presents antifouling characteristics. However, apart from the characterized potential increase in hydrophilicity to the plasma-treated SU-8 surfaces, further investigation on surface potential changes of the different samples are also necessary to check whether it plays a role in observed bacterial behavior. Although surface motility can be a crucial strategy in bacterial growth whereby bacteria seek nutrients and enable new colonies[39], the present results emphasize this mechanism by which the necessity of the affinity of pili to the substrate is required. Overall, the study provides comprehensive understanding of bacterial-SU-8 surface interaction which could lead to design effective strategies for the fabrication of SU-8 based various unique devices for biomedical applications[8–11,14].

## 4. Conclusion

In summary, our study elucidates the nanoscale surface changes of SU-8 thin films governed by wettability and surface carboxylic groups and how such properties influence bacterial motility, adhesion, and subsequent colonization in a spatiotemporal manner. The enhancement of the motility and direction of *X. fastidiosa* single cells on carboxylated surfaces during initial stages of growth has been observed. Particularly, the striking difference in cell trajectories and associated MSD values emphasize the underlying bacterial pili-surface interaction via surface functional groups. Notably, the hydrophobic nature of pristine SU-8 surfaces leads to relatively strong bacterial adhesion and larger number of microcolonies despite no observable motile cells. Conversely, the hydrophilic surface of plasma treated SU-8 samples exhibits significantly lower number of surface adhered single cells and about 70% lower bacterial coverage than the pristine SU-8 surfaces. Moreover, this study improves the current understanding of the role of surface functional groups and wettability on bacteria-surface interaction and thereby create strategies to prevent microbial adhesion and consequently, biofilm development of pathogenic species. Nevertheless, the present results warrant further biophysical investigation on how efficiently the different motility mechanisms can be deployed in view of type IV pili coordination and interaction with respect to surface properties changes. Further understanding of such bacterial-SU-8 interaction should lead to developing surfaces that can be tuned to adsorb or reject cell types, and thus design novel biological systems.

## ACKNOWLEDGEMENTS

The authors would like to thank Dr. Richard Janissen (TU Delft, Netherlands) for fruitful discussions. The authors are greatly indebted to Ursula F.S. Roggero (FEEC, UNICAMP) and Vitor Pelegati (IFGW, UNICAMP) for their assistance during sample preparation and confocal microscope access, respectively. This work was financially supported by the Brazilian agencies CNPq (441799/2016-7 and 429326/2018-1) and FAPESP (Grants number 2013/10957-0 and 2019/07616-3). S.A. and A.M.S acknowledge FAPESP (2017/14398-7) and CNPq, respectively, for funding their scholarships. The authors thank INFABIC/UNICAMP (FAPESP: 2014/50938-8, CNPq: 465699/2014-6), Semiconductor Component and Nanotechnology Center (CCSNano, UNICAMP) and the Device Research Laboratory (IFGW, UNICAMP) for granting access to their facilities.

