## Supplementary information for "Physiochemically distinct SU-8 surfaces tailor *Xylella fastidiosa* cell-surface holdfast and colonization"

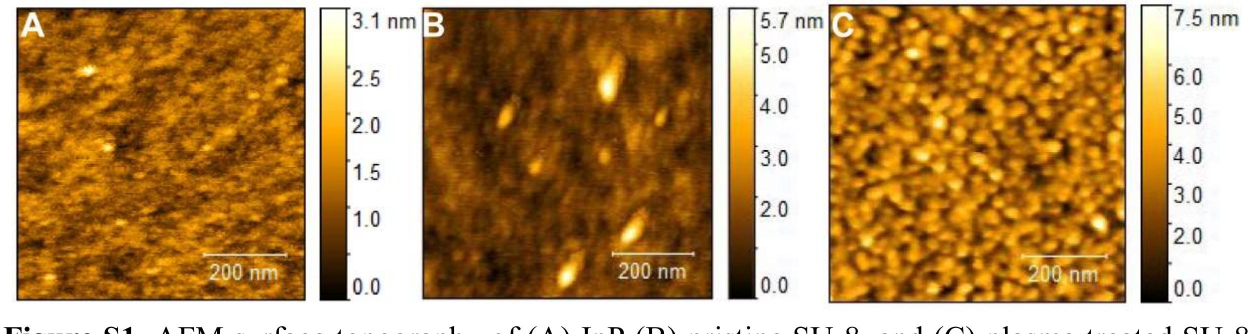


**Figure S1**. AFM surface topography of (A) InP (B) pristine SU-8, and (C) plasma treated SU-8 samples.

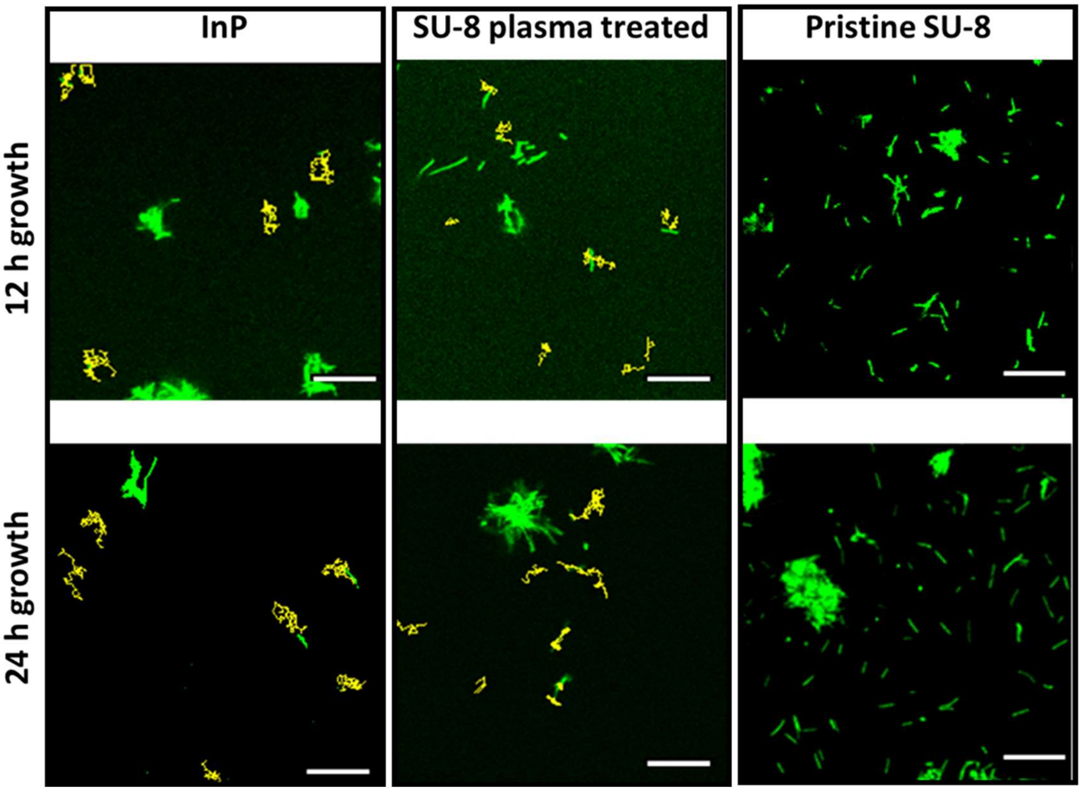


**Figure S2.** Trajectory of moving *X. fastidiosa* single cells on InP, SU-8 plasma-treated and pristine samples (no trackable moving cells) tracked from time lapse CLSM videos of 3 min after 12 and 24 h growth. Scale bar:20 µm

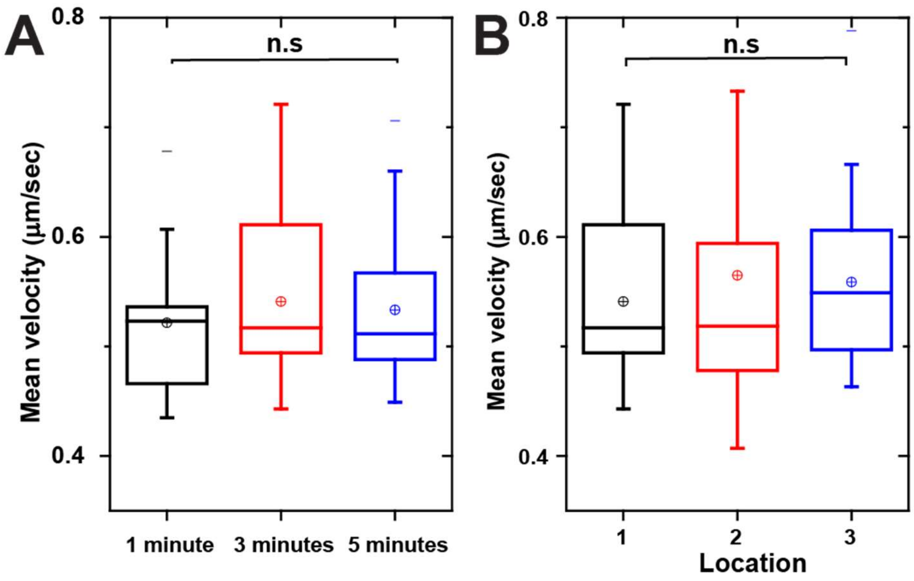


**Figure S3**. Mean velocity of single cell movement tracked for (A) 1, 2 and 3 mins, and (B) 3 different location for 3 mins of tracking on the plasma treated SU-8 surfaces after 6 h growth.

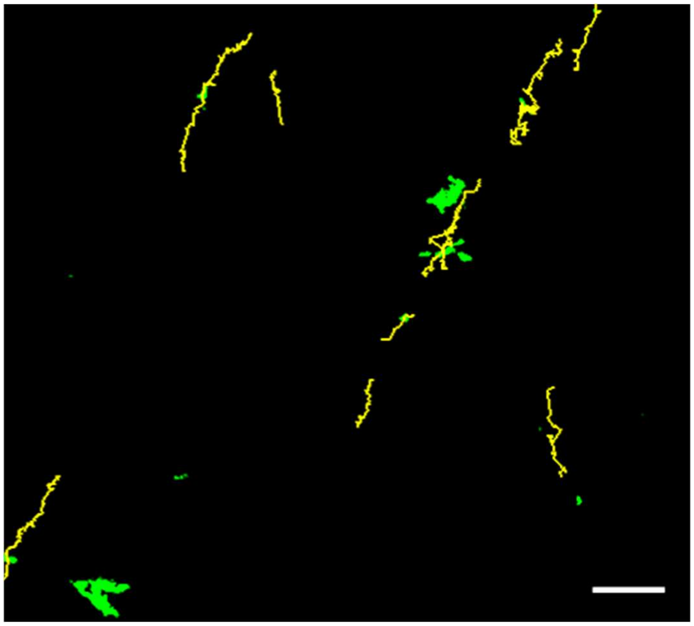

**Figure S4**. Trajectory of moving *X. fastidiosa* single cells on InP carboxylic functionalized samples from time lapse CLSM videos tracked for 3 minutes after 6 h growth. Yellow lines indicate cell trajectory. Scale bar:20 µm.

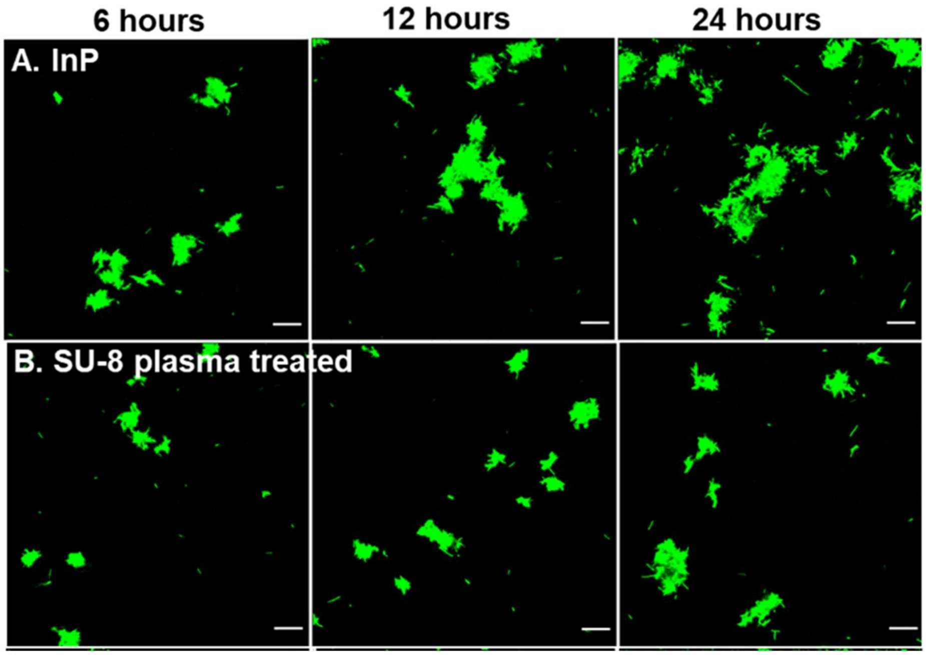


**Figure S5.** *In vitro* CLSM image of typical bacterial adhesion and microcolonies formation on surfaces of (A) InP and (B) plasma-treated SU-8, after 6, 12 and 24 hours of growth. Color histogram is adjusted so that single cells can be visualized. Scale bar:20 μm.

**Table S1.** Area (%) for the peaks fitted to angle distribution (negative and positive ranges) of individual segments of the cell trajectories for the plasma treated SU-8 surface for different growth times (from Figure 2E).

| **Growth time** | **Peak fit area (%)** | |
|---|---|---|
| | **Negative angle (-180$^o$ to 0$^o$)** | **Positive angle (0 to 180$^o$)** |
| **6 h** | 32.24 | 67.76 |
| **12 h** | 46.88 | 53.12 |
| **24 h** | 51.04 | 48.96 |
